# Single-Cell Analysis Reveals Regional Reprogramming during Adaptation to Massive Small Bowel Resection in Mice

**DOI:** 10.1101/615054

**Authors:** Kristen M. Seiler, Sarah E. Waye, Wenjun Kong, Kenji Kamimoto, Adam Bajinting, William H. Goo, Emily J. Onufer, Cathleen Courtney, Jun Guo, Brad W. Warner, Samantha A. Morris

## Abstract

**Background & Aims:** The small intestine (SI) displays regionality in nutrient and immunological function. Following SI tissue loss (as occurs in short gut syndrome, or SGS), remaining SI must compensate, or ‘adapt’; the capacity of SI epithelium to reprogram its regional identity has not been described. Here, we apply single-cell resolution analyses to characterize molecular changes underpinning adaptation to SGS.

**Methods:** Single-cell RNA-sequencing was performed on epithelial cells isolated from distal SI of mice following 50% proximal small bowel resection (SBR) vs. sham surgery. Single-cell profiles were clustered based on transcriptional similarity, reconstructing differentiation events from intestinal stem cells (ISCs) through to mature enterocytes. An unsupervised computational approach to score cell identity was used to quantify changes in regional (proximal vs distal) SI identity, validated using immunofluorescence, immunohistochemistry, qPCR, western blotting, and RNA-FISH.

**Results:** Uniform Manifold Approximation and Projection-based clustering and visualization revealed differentiation trajectories from ISCs to mature enterocytes in sham and SBR. Cell identity scoring demonstrated segregation of enterocytes by regional SI identity: SBR enterocytes assumed more mature proximal identities. This was associated with significant upregulation of lipid metabolism and oxidative stress gene expression, which was validated via orthogonal analyses. Observed upstream transcriptional changes suggest retinoid metabolism and proximal transcription factor *Creb3l3* drive proximalization of cell identity in response to SBR.

**Conclusions:** Adaptation to proximal SBR involves regional reprogramming of ileal enterocytes toward a proximal identity. Interventions bolstering the endogenous reprogramming capacity of SI enterocytes—conceivably by engaging the retinoid metabolism pathway—merit further investigation, as they may increase enteral feeding tolerance, and obviate intestinal failure, in SGS.

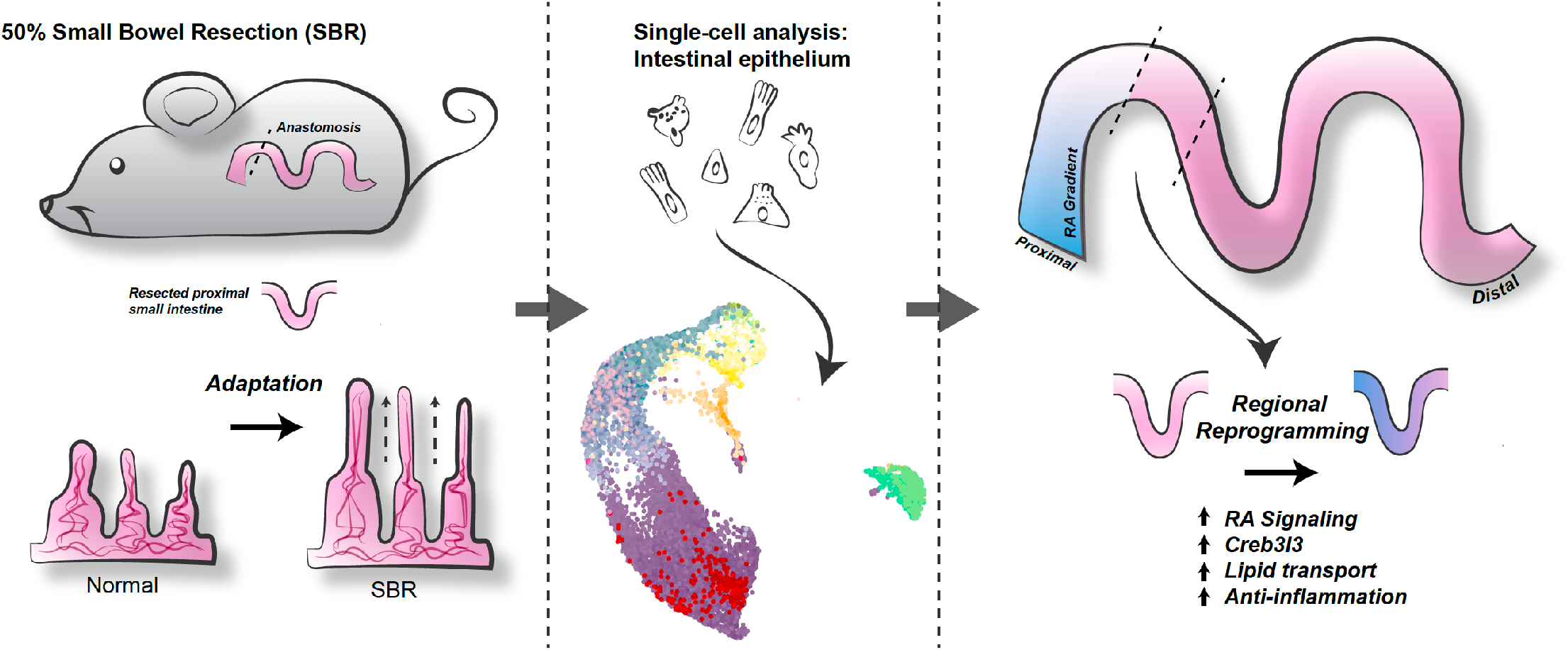

**Synopsis:** Here, single-cell RNA sequencing reveals interactions between the retinoid metabolism pathway and ‘regional reprogramming’ of distal small intestinal epithelium to a proximal identity following proximal small bowel resection. This provides novel insight into physiological adaptation to short gut syndrome.

## Background

The small intestine (SI) absorbs nutrients necessary to sustain life, and displays regional specialization for absorption of specific nutrients along its cephalocaudal axis from duodenum to jejunum to ileum. The majority of nutrient absorption occurs in the duodenum and jejunum, while the ileum is primarily responsible for absorbing bile, Vitamin B12, and fat-soluble vitamins. The ileum is also more prone to inflammatory disorders relative to proximal intestine, owing in part to higher bacterial load.

A variety of diseases require surgical resection of significant lengths of SI. These range from congenital anomalies and necrotizing enterocolitis in children to trauma, embolism, and malignancy in adults. The resulting loss of SI can cause short gut syndrome (SGS), or the inability of the SI to completely support the metabolic demands of a patient. Management options for SGS are limited, comprising parenteral nutrition (PN), intestinal lengthening procedures, and ultimately small bowel transplant, all incurring significant morbidity and mortality^1, 2^.

Our murine model of SGS is based on small bowel resection (SBR), where 50% of the proximal SI is surgically removed^3^. Sham surgery consists of transection and anastomosis, without removal of SI, and acts as a control for exposure to anesthesia, laparotomy, and intestinal transection. This model elicits villus lengthening in the remnant ileum of SBR but not sham mice,^3^ yielding increased mucosal absorptive surface area to compensate for lost tissue. Importantly, the degree of this “structural adaptation” response correlates with “functional adaptation,” as evidenced by increased oral tolerance and weight gain in mice. This model is relevant to clinical SGS, as structural adaptation correlates with oral tolerance and weaning from PN observed in human patients^4^. At the same time, structural adaptation does not intrinsically predict functional adaptation, as perturbed weight gain and steatorrhea affect mice deficient in CXCL5 after SBR, despite normal structural adaptation^5^. This leads us to conclude that additional factors beyond simple tissue hyperplasia are at play, which are likely underscored by molecular changes at the single-cell level.

While structural and, to a lesser extent, functional adaptation following SBR have been characterized, relatively little is understood about the molecular changes that accompany the adaptation process. In this respect, mRNA- and protein-level expression analyses offer crucial insight, as clinically appreciable adaptation may require that cells assume molecular identities mimicking those of the resected region. Studies of adaptation at the structural and functional level lack the resolution to explore this possibility. Here, in the case of proximal SBR, we hypothesize that remnant ileum (distal SI) upregulates gene and protein expression patterns characteristic of the jejunum (proximal SI) at the single-cell level, a process we term “regional reprogramming”.

Clinical therapies to induce regional reprogramming could enhance an SGS patient’s ability to tolerate oral intake and wean from PN by augmenting the innate functionality of epithelial cells. It is possible this approach may actually be *more* effective than the previous “holy grail” of SGS research, which has primarily focused on inducing structural adaptation. Enhanced structural adaptation is intrinsically more metabolically demanding (tissue growth) and may or may not affect the key absorptive, metabolic, and immunological pathways specifically deficient in a SGS patient.

To test our hypothesis and address gaps in our understanding of adaptation to SGS, we employed high-throughput single-cell RNA sequencing (scRNA-seq) to characterize gene expression changes of distal SI epithelium during adaptation following massive proximal SBR. This allowed us to dissect population heterogeneity within the epithelium, and to characterize the regionalization pathways critical in the adaptive response. Here, we show that following SBR, the SI epithelium ‘regionally reprograms’ toward mature proximal enterocyte identity, accompanied by increased proximal SI nutrient processing gene expression. These changes are punctuated by the increased expression of the proximal SI transcription factor, *Creb3l3*, a key candidate for reprogramming distal SI gene regulatory networks to a more proximal identity. Analysis of upstream pathways suggests a role for retinoic acid (RA) signaling in driving the adaptation response. Together, our single-cell analyses have enabled high-resolution characterization of the molecular changes and regional reprogramming that underlie adaptation.

## Results

### Cellular heterogeneity of the small intestinal epithelium is captured by scRNA-seq

Structural adaptation reliably occurs by day 7 after SBR. As such, at day 7 after sham or SBR surgery, typical SBR structural adaptation was confirmed, with villi height increasing by 86.19 ± 19.14 μm (*p* < 0.01) relative to sham (**Figure 1a**). Epithelial cells were harvested from SI in an area equidistant from the anastomosis, dissociated into single cells, and processed via high-throughput droplet-based scRNA-seq using the 10x Genomics platform^6^. In total, we sequenced 19,245 cells from 9 independent biological replicates (Sham: 8,209 cells, n=5 replicates; SBR: 11,036 cells, n=4 replicates). A mean of 1,767 and 1,763 genes per cell in sham and SBR, respectively, and 6,754 and 6,111 transcripts per cell in sham and SBR, respectively, were detected (**Figure 1b**).

**Figure 1.**
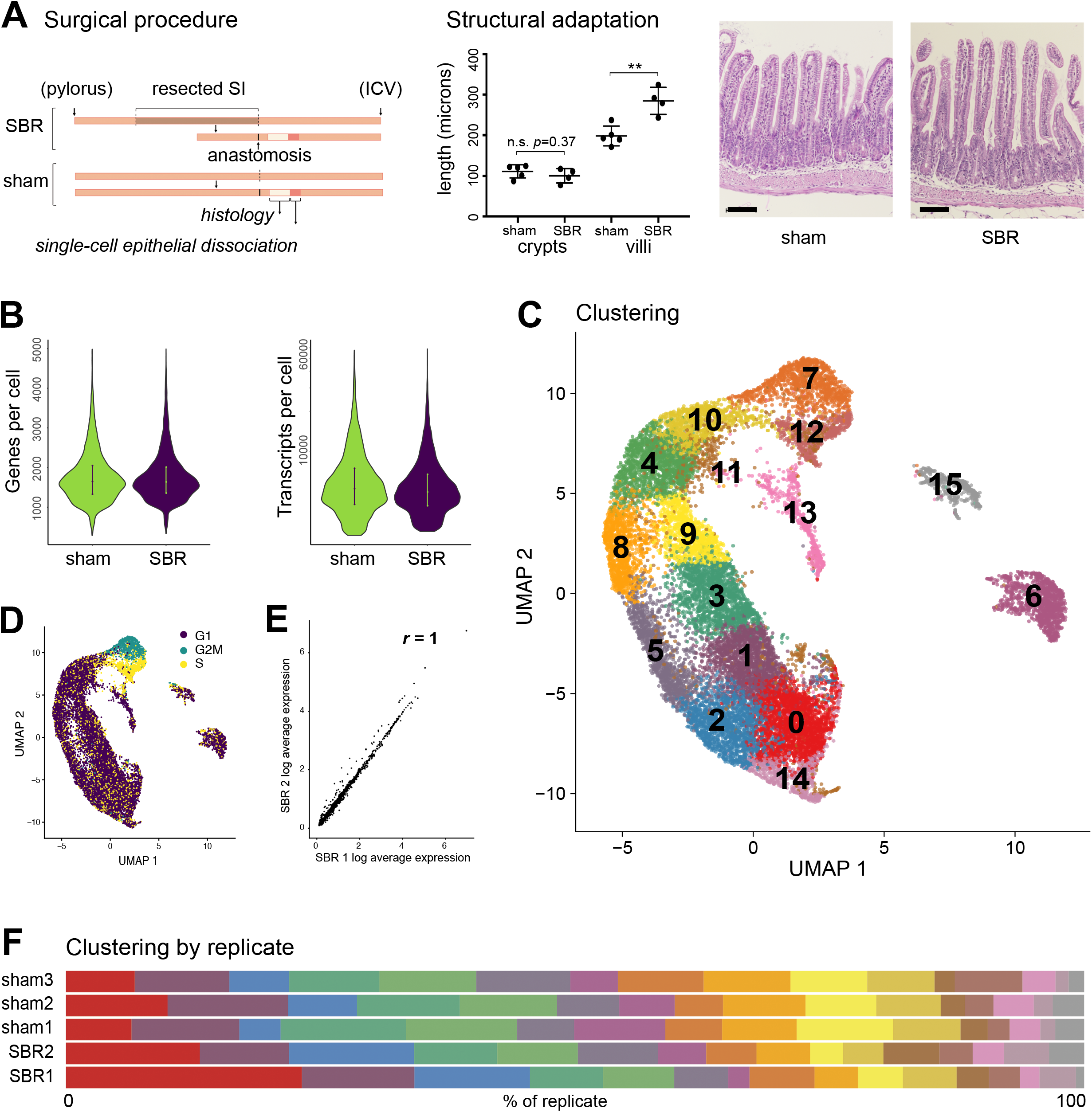
Experimental design, quality control, and single-cell analysis. **A)** 50% proximal small bowel resection (SBR) and sham operation were performed on mice. 7 days after surgery, the intestine distal to the anastomosis (ileum) was harvested and equal amounts of tissue equidistant from the anastomosis were used to generate single cell epithelial suspensions. An area immediately adjacent to this was prepared for histological examination. Typical structural adaptation of SBR mice (lengthened villi) was confirmed (*p*=0.003), with a representative hematoxylin and eosin image of SI tissue from a sham vs SBR mouse shown (20X image acquired using Nikon Eclipse 80i, scale bar= 100 μm). Epithelium from mice demonstrating structural adaptation was prepared for scRNA-seq analysis. **B)** A mean of 1,767 and 1,763 genes per cell in sham and SBR, respectively, and 6,754 and 6,111 transcripts per cell in sham and SBR, respectively, were detected. **C)** Uniform Manifold Approximation and Projection (UMAP) of integrated biological replicates identified 16 unique cell clusters. **D)** Cell cycle states projected onto the UMAP. **E)** Representative plot of SBR experimental replicates demonstrated similar gene expression profiles. Correlation coefficient (R) of average gene expression are as shown between these biological replicates. Total biological replicates were n=5 sham and n=4 SBR (n=3 “sham1”, n=1 “sham2”, n=1 “sham3”, n=3 “SBR1”, n=1 “SBR2”). **F)** The same 16 clusters were identified in both sham and SBR, in all replicates. Distribution of all cells across clusters 0 through 15 (from left to right), by replicate, is shown.

To cluster and visualize cells based on their transcriptional similarity, we used the R package, Seurat^7, 8^, and Uniform Manifold Approximation and Projection (UMAP)^9^. UMAP analysis and plotting revealed 16 clusters of transcriptionally distinct cell types/states (**Figure 1c)**, where clustering was not driven by numbers of detected genes or transcripts (not shown). Scoring and projection of cell cycle state onto the UMAP plot revealed clustering of cells in S and G1 phases, corresponding to stem cells and transit amplifying (TA) cells (**Figure 1d**; and below). Gene expression between equivalent biological replicates was highly correlated, demonstrating a high degree of consistency between the independent biological replicates (**Figure 1e**). Furthermore, cells from every cluster were represented in each biological replicate, demonstrating consistency of cell capture (**Figure 1f**).

To assign cell identity to each cluster in an unsupervised manner, we used a computational method based on quadratic programming (QP)^10^ to score individual cell identity against an existing single-cell atlas of well-annotated SI cell types^11^. This reference atlas contains stem cells, TA cells, early and late enterocyte progenitors, immature proximal and distal enterocytes, mature proximal and distal enterocytes, goblet cells, Paneth cells, enteroendocrine cells, and tuft cells, annotated based on an extended list of previously identified markers^11^. Scoring cell identity using QP is beneficial because it can capture cells in transitional states, rather than assigning cell identities as binary values. This is ideal for assessing developmental and disease processes where cell identity can be considered as continuous rather than discrete. This flexibility is not found in most classification systems that definitively assign discrete identities to cells, without considering their transitional states. We have previously demonstrated the efficacy of QP in placing cells into an identity continuum during lineage reprogramming^12^. Considering enterocytes progressively transdifferentiate while migrating along the villus axis^13^, and our hypothesis that cell identity is reprogrammed during adaptation, QP represents an appropriate method for quantifying any changes in cell identity that accompany SBR.

Cell identity scores generated by QP were projected onto the UMAP plot, enabling cell cluster identity to be annotated (**Figure 2a**). This confirmed that all major cell types of the SI epithelium, as above, were captured by our single-cell analysis, including low numbers of tuft cells and enteroendocrine cells (0.3% and 0.03%, respectively, as a proportion of all captured cells). Projection of all identity scores onto the UMAP plot demonstrated this clustering and visualization method does indeed retain both local and global information, capturing the differentiation trajectory from stem cells to mature enterocytes (**Figure 2b**). Further examination of these clusters revealed that expression of the proliferation marker *Mki67* is enriched in stem, TA, and progenitor cells, and is downregulated as cells begin to differentiate. Conversely, expression of mature enterocyte marker alkaline phosphatase (*Alpi*) increases in concert with differentiation/maturation, with highest expression colocalizing in areas identified by QP as mature enterocytes (**Figure 2c**). Furthermore, based on this differentiation trajectory, we posited the most terminal cluster on the UMAP plot would be enriched for markers specific to villus tip cells, such as adenosine deaminase (*Ada*)^13^, which we confirmed (**Figure 2c**). Finally, one population (cluster 15) did not receive a definitive QP score for any of the reported SI epithelial lineages^11^. This cluster is enriched for intraepithelial lymphocyte (IEL) marker expression, including *Cd45* (**Figure 2c**), suggesting these cells represent IELs captured alongside the SI epithelium.

**Figure 2.**
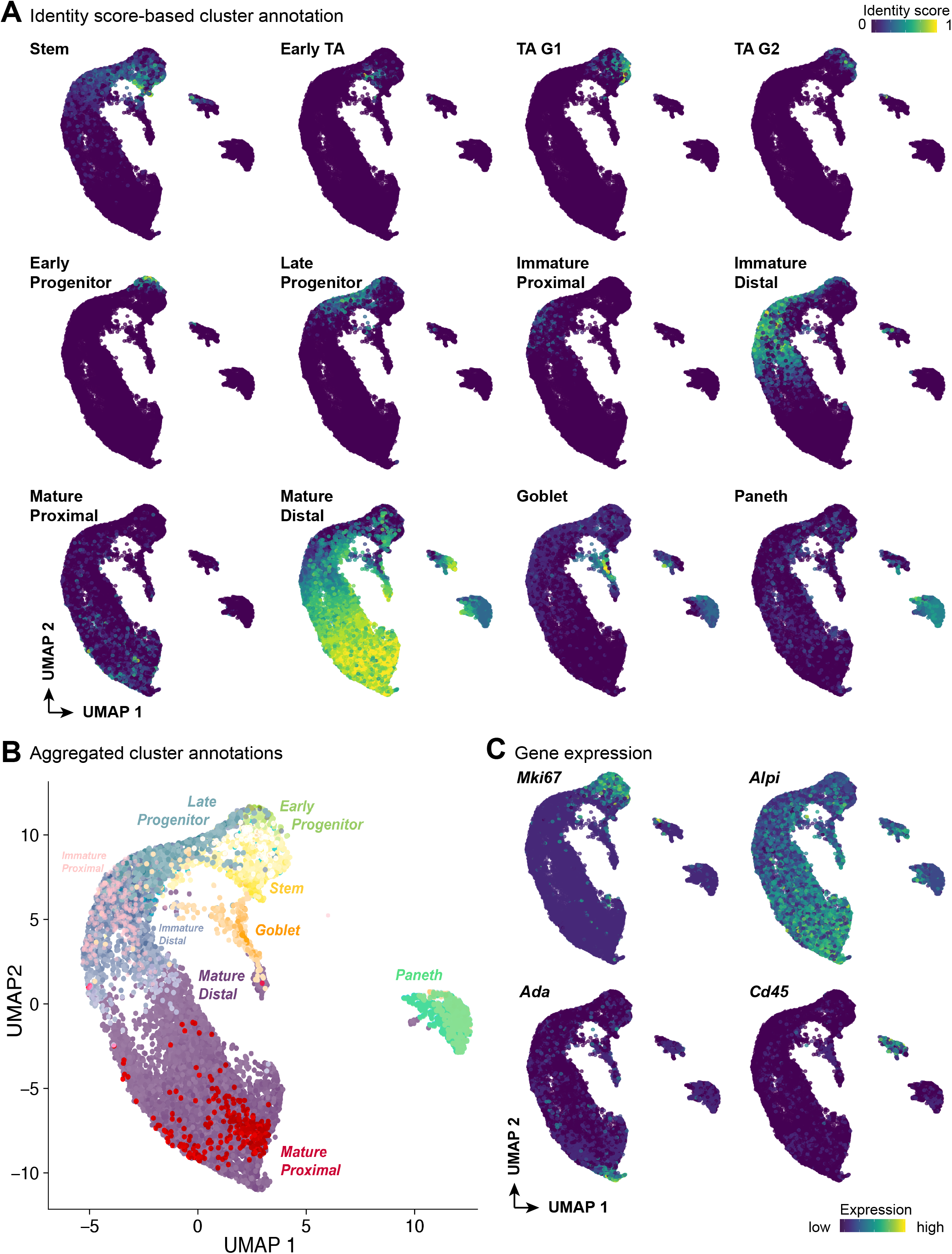
Annotation of cell identities using Quadratic Programming (QP). **A)** QP-based identity scores of intestinal epithelial populations, projected onto UMAP. Cell populations (from left to right) include stem, early transit amplifying (TA), TA G1, TA G2, early enterocyte progenitors, late enterocyte progenitors, immature proximal enterocytes, immature distal enterocytes, mature proximal enterocytes, mature distal enterocytes, goblet cells, and Paneth cells. Low percentages of tuft and enteroendocrine cells (0.3% and 0.003%, respectively) were identified and are not shown. **B)** Aggregated QP scores provide a summary of cell identities within the UMAP, demonstrating a maturation trajectory from stem cells to mature enterocytes. **C)** Projection of transcript enrichment for selected QP validation markers, clockwise from top left: Proliferation marker Antigen KI-67 (*Mki67*) is appropriately enriched in the stem, TA, and progenitor regions of UMAP; Alkaline phosphatase (*Alpi*) expression increases as enterocyte maturation occurs; *Cd45* identifies a population enriched for intraepithelial lymphocytes (IELs); Adenosine deaminase (*Ada*), a villus tip marker, localizes at the termination point of the developmental trajectory. Color scale bar indicates relative intensity of cell identity scoring (A) or gene expression (C) across the UMAP.

In summary, our single-cell analyses identified all major SI epithelial cell populations, again demonstrating the strength of our unsupervised QP cell classification approach in concert with UMAP visualization to identify cells differentiating along a defined trajectory. Together, this provides a comprehensive picture of SI epithelial heterogeneity, under both sham and SBR conditions.

### Quantification of SI epithelial cell composition changes following SBR reveals ‘regional reprogramming’ toward mature proximal enterocyte identity

Changes in cell type composition of the SI epithelium accompany adaptation. However, reports on the precise nature of these changes have been conflicting, with studies describing a relative expansion of either enterocytes or secretory lineages following SBR^14–18^. Discrepancies between these studies may arise due to differences in resection location, amount, the use of differing animal models, and experiment conditions or durations. Furthermore, these previous studies were limited by a lack of resolution to assess cell identity in an unbiased manner. Here, we use our unbiased single-cell resolution classification of cell identity to precisely quantify changes in epithelial composition following SBR. We assessed the distribution of sham- vs SBR-derived cells by projecting the densities onto the UMAP plot (**Figure 3a)**. First, looking at the villus enterocyte population (comprised by immature and mature enterocytes), we found a significant increase following SBR, as a total of all cells surveyed (68.9% ± 3.1% of sham events sampled, vs 76.8% ± 0.1% of SBR (*p*<0.05); **Figure 3b**). This increase in villus enterocytes, at single-cell resolution, is in agreement with immunohistochemistry analysis of our tissue samples. Here, we identified villus enterocytes as non-goblet (mucin-2, or MUC2 expressing) nucleated villus cells divided by all nucleated cells of the crypt-villus axis, revealing that enterocytes comprise 71.6% of sham epithelium, vs 74.9 % of SBR epithelium (**Figure 3c**). From our single-cell analysis, we did not observe significant changes (*p*=0.06) in the proportion of secretory lineages comprising sham (22.3% ± 5.8%) and SBR (13% ± 0.3%) SI epithelium (**Figure 3b**).

**Figure 3.**
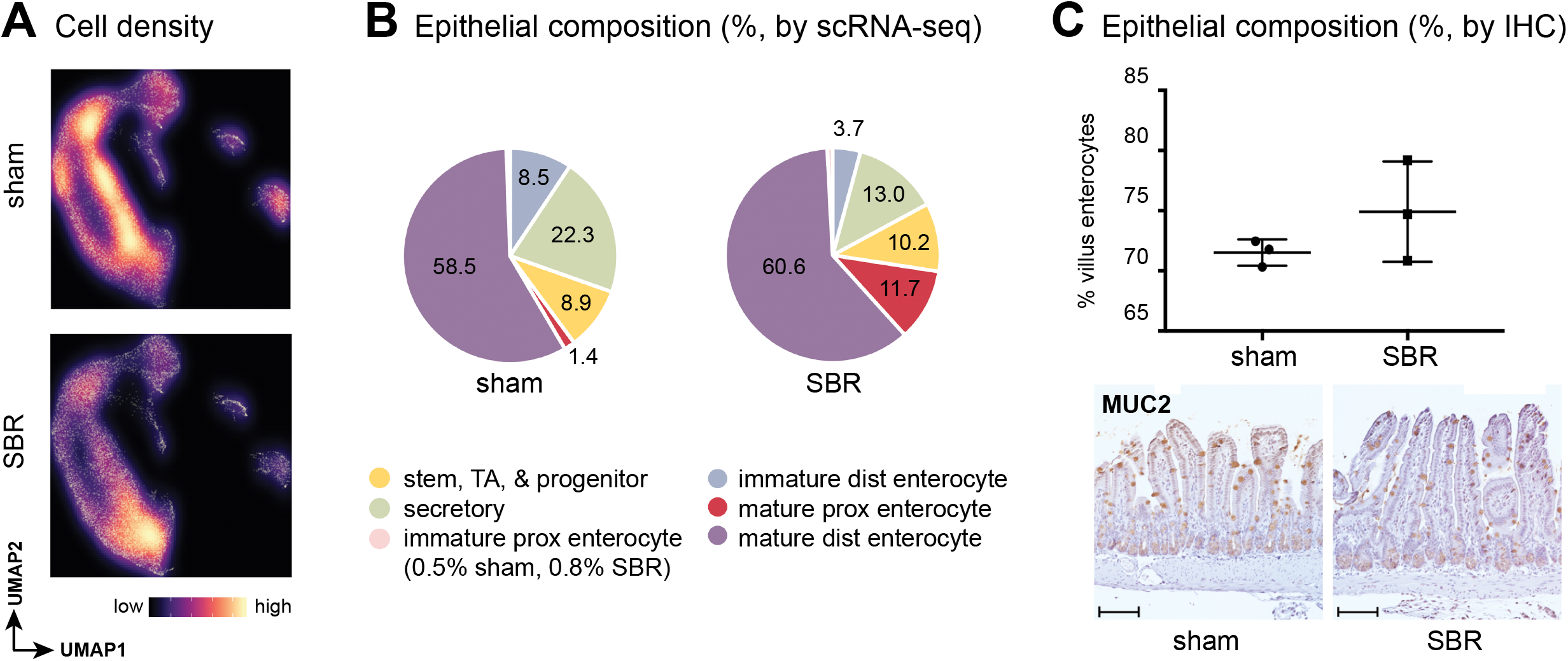
Relative expansion of mature proximal enterocytes occurs in SBR mice. **A**) Density of sampled cells from sham and SBR epithelium, projected onto the UMAP, demonstrates a population shift toward mature proximal enterocytes in SBR. Color scale bar indicates relative density. **B**) Graphical representation of how epithelial lineages (as identified by QP), contribute to the total composition of events sampled by scRNA-seq in sham vs SBR. **C**). Immunohistochemical (IHC) analysis was performed on tissue samples (n=3 sham and n=3 SBR) to confirm the relative expansion in villus enterocytes (i.e. immature and mature enterocytes) as a percent of total epithelium in SBR, as predicted by scRNA-seq. Representative images of IHC for mucin 2 (*Muc2*, goblet cell marker) in sham vs SBR are provided (20X image acquired using Nikon Eclipse 80i, scale bar= 100 μm).

Continuing to focus on the enterocyte lineage, as this absorptive cell type is central to adaptation, we next quantified enterocyte differentiation between sham and SBR populations. We found that sham samples contained 9.0% ± 5.5% immature and 59.9% ± 7.2% mature enterocytes, compared to 4.5% ± 2.5% immature and 72.3 ± 2.7% mature in SBR (**Figure 3b**). Though these differences in immature and mature enterocyte composition between sham and SBR did not reach significance (*p*=0.2 and 0.06, respectively), this observation is supported by a previous report of enhanced metabolically mature enterocyte migration after SBR^19^.

Considering our hypothesis that cells adopt a different regional identity to aid adaptation, we next quantified the balance between proximal and distal enterocyte identities in sham vs. SBR samples. As expected considering the tissue was harvested from ileum, a large percentage of cells (58.5% ± 7.5%) from sham samples received high mature distal enterocyte scores with very few cells (1.4% ± 0.6%) scoring as mature proximal enterocytes (**Figure 3b**). In contrast, in SBR samples we found a significant increase in the percentage of cells receiving high mature proximal enterocyte scores (11.7% ± 4.1%, *p*<0.05), (**Fig. 3b**). This shift toward mature proximal enterocyte identity in SBR suggests a transcriptional “proximalization,” or regional reprogramming, of ileum in response to proximal SI resection.

### Regional reprogramming of distal SI after SBR is accompanied by increased proximal SI nutrient processing gene expression

From our above analyses, changes in SI epithelial composition following SBR center primarily on a shift toward mature proximal enterocyte identities, suggesting these changes are a key driver of the adaptive response. Thus, we next focused on characterizing the transcriptional changes underlying the shift toward mature proximal enterocyte identity in SBR.

To identify significant transcriptional changes after SBR, we performed differential gene expression analysis, identifying 174 differentially expressed genes between all sham and SBR cells. The 10 most significantly expressed genes in SBR, relative to sham, are shown in **Table 1**. This list of SBR-associated transcripts is enriched for signature genes of proximal SI nutrient processing function, including apolipoprotein A-IV (*Apoa4*), fatty acid binding protein 1 (*Fabp1*), apolipoprotein C-III (*Apoc3*), lactase (*Lct*), and epoxide hydrolase 2 (Ephx2)^11, 20^ (**Figure 4a**). In contrast, distal SI transcript fatty acid binding protein 6 (*Fabp6*)^11^ was significantly depleted in SBR (0.62 average log fold change depleted, *p*<0.001) (**Fig. 4a**).

**Figure 4.**
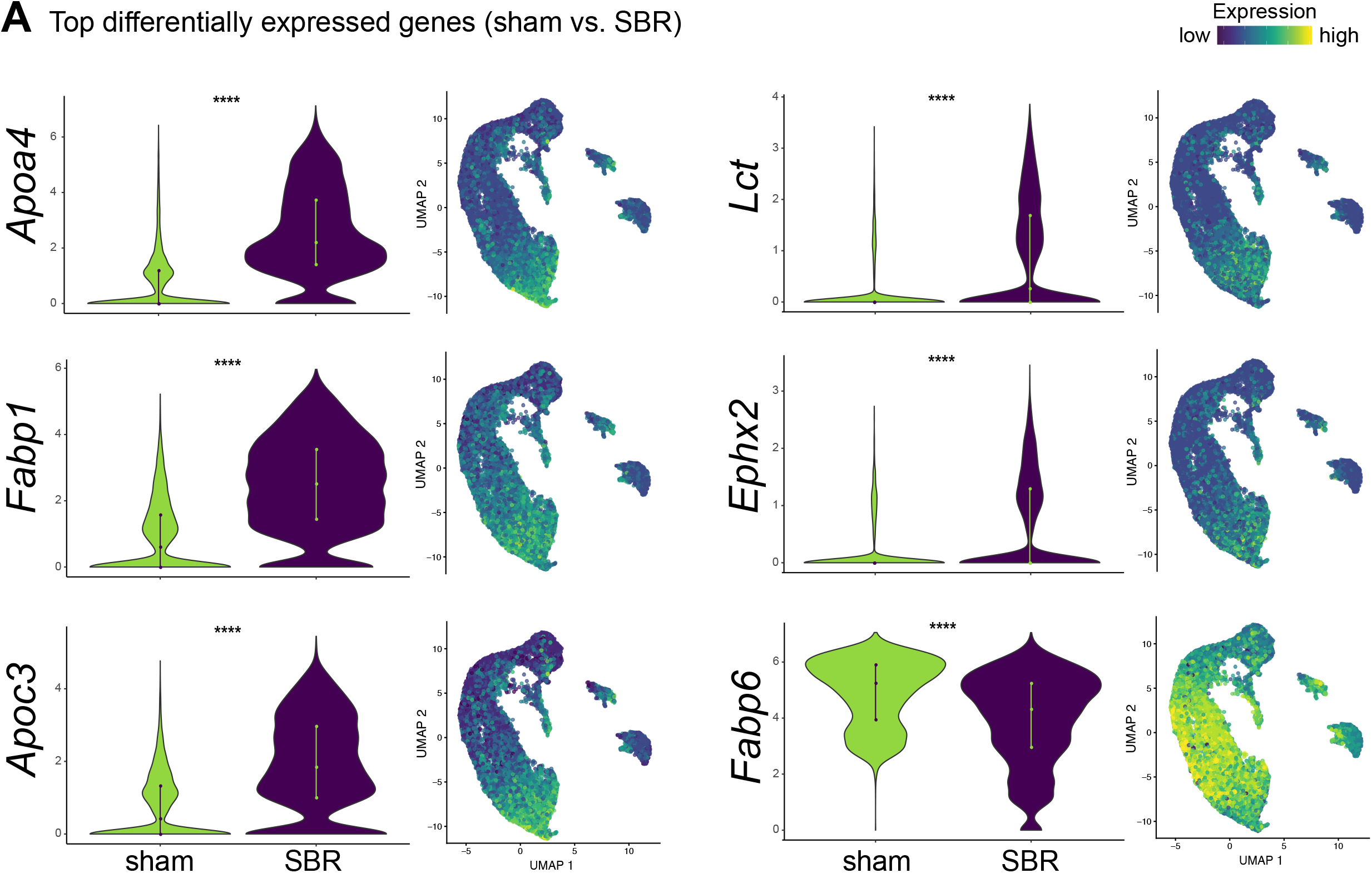
Identification of signature proximal small intestine transcripts that increase after SBR. **A)** Violin plots (left) show relative expression of transcripts in sham vs, SBR populations, and UMAP plots (right) show relative transcript expression levels within cell populations.

**Table 1:** Top 10 genes upregulated in SBR relative to sham epithelium, with average log fold change (logFC) and adjusted *p-*values (p val)

| Gene | logFC | p val | Gene | logFC | p val |
| --- | --- | --- | --- | --- | --- |
| *Apoa4* | 2.05 | <0.001 | *Fabp2* | 0.96 | <0.001 |
| *Fabp1* | 1.76 | <0.001 | *Sepp1* | 0.92 | <0.001 |
| *Apoc3* | 1.46 | <0.001 | *Leap2* | 0.79 | <0.001 |
| *Lct* | 1.11 | <0.001 | *Ephx2* | 0.71 | <0.001 |
| *Rbp2* | 1.10 | <0.001 | *Apob* | 0.69 | <0.001 |

Increased *Apoa4* and *Rbp2* expression following SBR has been previously described^21–23^. Contrary to our findings, one of these studies reported no significant changes in ileal *Fabp1* mRNA expression. However, this study examined whole tissue preparations, rather than epithelium at single-cell resolution^23^. Thus, to validate the transcriptional changes revealed by our scRNA-seq analysis, we surveyed *Fabp1* expression via RNA-FISH on histological sections of sham and SBR animals at 7- and 70-days post-surgery, with the latter analysis designed to investigate whether the observed changes are stable, a property rarely investigated in the context of SBR. At day 7, *Fabp1* showed a 1.5 average log2 fold A.U. (Arbitrary Units) increased expression per nucleated villus cell in SBR, relative to sham (*p*<0.001), and this response was maintained through day 70 post-surgery (1.4 average log2 fold A.U. increased expression, *p*<0.01) (**Figure 5a**).

**Figure 5.**
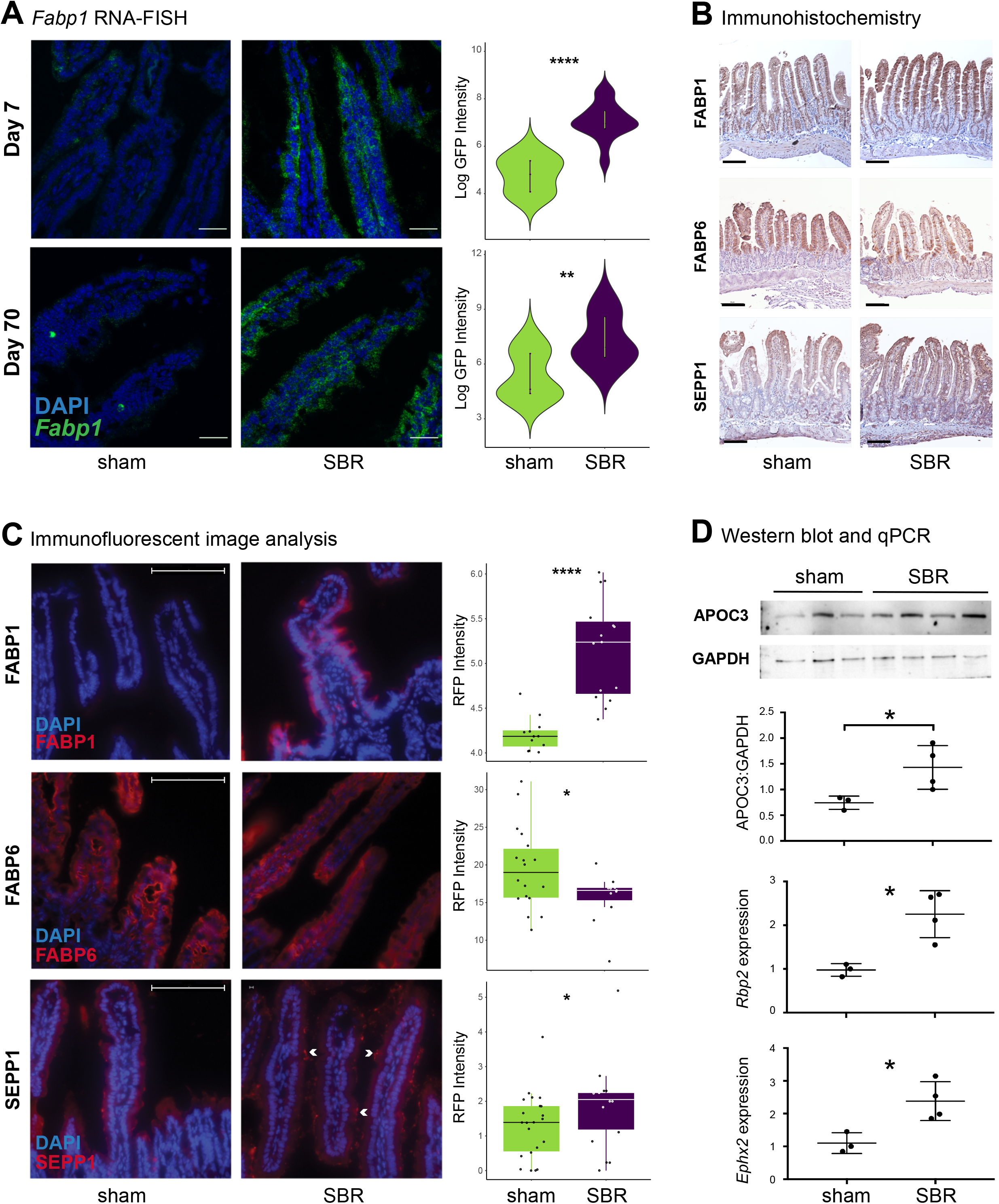
Validation of proximal small intestine markers that increase after SBR. **A)** RNA fluorescence in situ hybridization (FISH) for *Fabp1* shows significant upregulation of transcripts (GFP) per nucleated cell (DAPI) at days 7 and 70 after surgery. Day 7: n=15 sham images, n=15 SBR images (3 biological replicates). Day 70: n=12 sham images, n =15 SBR images (3 biological replicates) (images acquired using Olympus FV1200 Confocal Microscope). **B)** Immunohistochemistry (IHC) staining for FABP1, FABP6, and SEPP1 in sham and SBR mice at post-operative day 7 shows qualitative increases in FABP1 and SEPP1, and decrease in FABP6, in SBR mice (images acquired using Nikon Eclipse 80i with Ds-Ri2 camera). **C)** Immunofluorescence staining images for FABP1 (n=12 sham images, n=15 SBR images, 3 biological replicates), FABP6 (n=18 sham images (4 biological replicates), n =11 SBR images (3 biological replicates)), and SEPP1 (n=22 sham images (5 biological replicates), n=18 SBR images (4 biological replicates)) were computationally analyzed to confirm significant protein-level changes corresponding to mRNA changes. White arrows indicate areas of intense SEPP1 expression in SBR (images acquired using Nikon Eclipse 80i with Ds-Ri2 camera). **D)** Top to bottom: Western blot analysis of APOC3 in sham and SBR epithelial lysates, normalized to GAPDH and quantified (n=3 sham and n=4 SBR mice); qPCR validation of upregulated SBR genes *Rbp2* and *Ephx2* from SI tissue (n=3 sham and n=4 SBR mice). RNA-FISH images are at magnification 60X, scale bar = 30 μm. IHC are at 20X, scale bar= 100 μm. IF are at 40X, scale bar= 100 μm. All graphs are presented as mean ± SD. *= *p*<0.05, **= *p*<0.01, ****= *p*<0.0001.

To further validate our findings, we performed quantitative immunofluorescence, western blotting, and qPCR for selected proteins and transcripts, including FABP1, FABP6, APOC3, *Rbp2*, and *Ephx2*. Representative immunohistochemistry images of FABP1 and FABP6 show qualitative changes in these proteins consistent with our scRNA-seq analysis (**Figure 5b**). Quantification of this immunostaining panel using immunofluorescence demonstrated significant increases in FABP1 (1.2 average log2 fold A.U., *p*<0.001) in SBR relative to sham (**Figure 5c**). In contrast, FABP6 was significantly decreased in SBR compared to sham (−1.2 fold relative fluorescent intensity, *p*<0.05) (**Fig. 5c**). Western blotting and quantification of APOC3 showed a 1.92 fold increase in SBR compared to sham (*p*<0.05), and qPCR confirmed up-regulation of *Rbp2* and *Ephx2* (**Figure 5d**).

*Sepp1* upregulation during adaptation to SBR is a novel finding, warranting further investigation. SEPP1 is a secreted glycoprotein with important immunomodulatory and antioxidant effects in the intestine^24–26^. Representative immunohistochemistry images of SEPP1 show qualitative changes consistent with our scRNA-seq analysis (**Figure 5b**). Quantification of this immunostaining via immunofluorescence demonstrated significant increases in SEPP1 (1.4 fold relative fluorescent intensity, *p*<0.05) in SBR relative to sham (**Figure 5c**). Furthermore, SEPP1 has been shown to suppress inflammation-associated tumorigenesis, in part through its effects on macrophage polarization, more specifically, by suppressing M2 associated *Ym1* expression^25, 27^. Interestingly, this is consistent with RNA-sequencing analysis we performed on sub-epithelial tissue from sham and SBR mice, showing that *Ym1* is the second most depleted transcript in SBR mice (−6.19 fold depleted, *p* <0.05, unpublished). This suggests a role for SEPP1 in mitigating oxidative-stress induced injury during adaptation (which is known to occur after SBR^18^, consistent with the fact that FABP1 directs fatty acids toward oxidative metabolism^28^), possibly via immunoregulatory effects on macrophages, and merits further study.

Together, these orthogonal validations confirm our scRNA-seq results, demonstrating that gene expression programs to support proximal SI nutrient processing, and to counteract the associated oxidative stress, are engaged following SBR. These mRNA- and protein-level changes underlie the regional reprogramming to mature proximal identity we observe in SBR.

### Proximal SI transcription factor *Creb3l3* shows stable upregulation accompanied by expanded villus axis zonation in SBR mice

We next aimed to identify candidate transcription factors responsible for driving the observed regional reprogramming and shift toward a mature proximal nutrient processing profile following SBR. Of 44 transcription factors previously shown to be differentially expressed between proximal vs distal enterocytes^11^, only one associated with proximal identity, cAMP responsive element binding protein 3 like 3 (*Creb3l3*), was upregulated in SBR (0.46 average log fold increase in SBR, *p*<0.0001). *Creb3l3* is a master regulator of lipid metabolism^29^, fitting with increased lipid metabolism after SBR. An additional transcription factor, Kruppel-like factor 4 (*Klf4*) was also increased in SBR (0.26 average log fold increase in SBR, *p*<0.001). Though *Klf4* was identified as a distal enterocyte transcription factor by Haber *et al*. ^11^, another group reported highest *Klf4* expression in duodenum and jejunum^30^, and described that *Klf4* plays a role in the maturation of intestinal stem cells, as well as in the differentiation of absorptive lineages, such as enterocytes^30^. It is worth noting the possibility that a wider variety of proximal transcription factors were upregulated immediately after surgery and stabilized to baseline by day 7, when structural adaptation was complete. However, since these were the only previously identified^11^ regional enterocyte transcription factors significantly differentially expressed in SBR at day 7, we focused our analysis on these factors as putative drivers of stable regional reprogramming.

First, we sought to determine whether proximal SI transcription factor *Creb3l3* expression was transiently upregulated in SBR, or whether it was critical to maintaining a long-term adaptive response. Since *Creb3l3* regulates lipid metabolism, and the increased demand for ileal lipid absorption should persist indefinitely after SBR, we expected its expression to remain elevated. Indeed, RNA-FISH for *Creb3l3* at days 7 and 70 demonstrated its significant and long-term upregulation following SBR (Day 7: 1.3 average log2 fold A.U. increase in SBR, *p*<0.001; Day 70: 1.2 average log2 fold A.U. increase in SBR, *p*<0.01) (**Figure 6a**).

**Figure 6.**
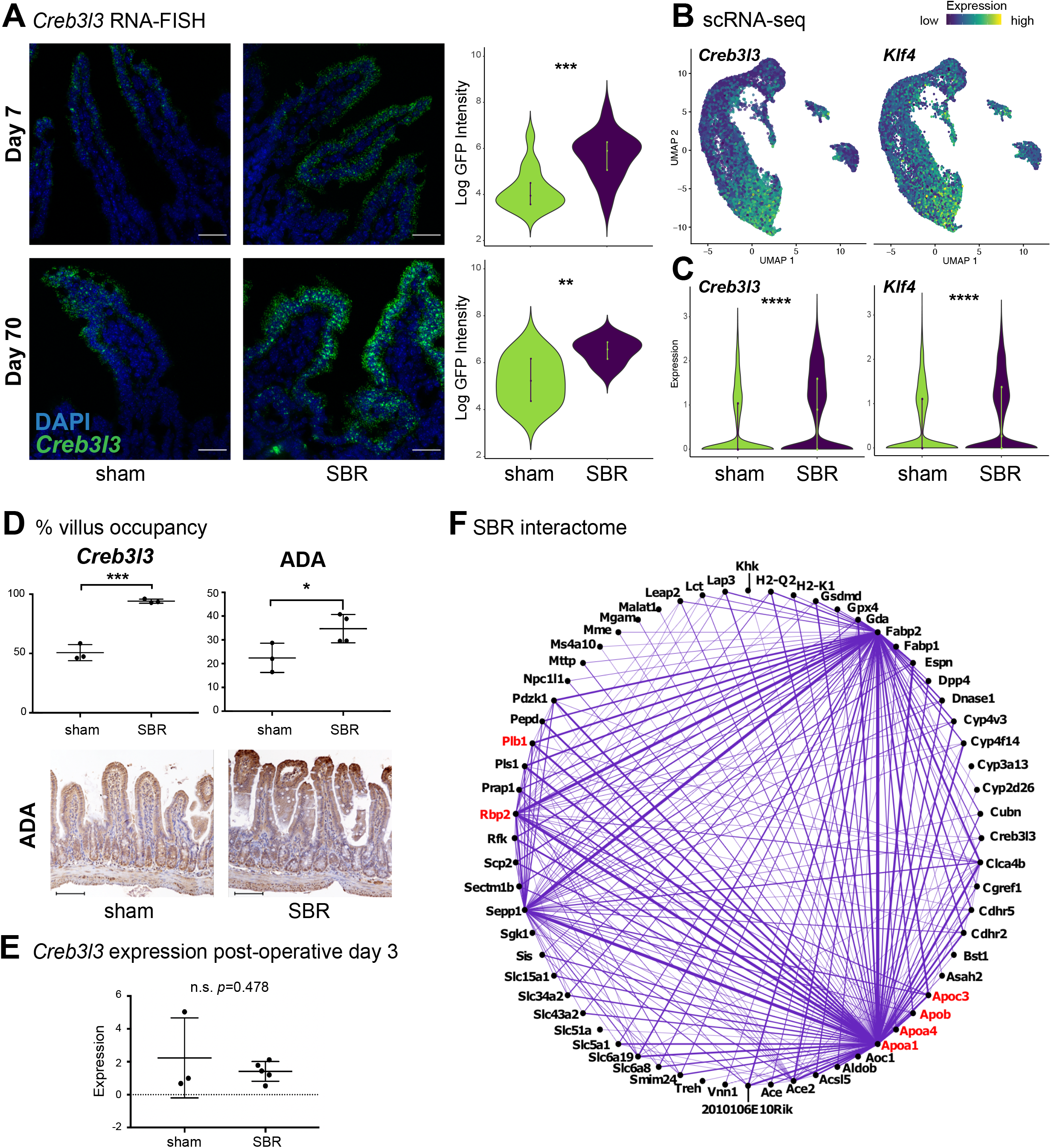
Dissecting genetic underpinnings of epithelial proximalization following SBR. **A)** RNA- FISH for *Creb3l3* shows significant upregulation of transcripts (GFP) per nucleated cell (DAPI) at days 7 and 70 after surgery. Images are at 60X, scale bar = 30µm, acquired using Olympus FV1200 Confocal Microscope. Day 7: n=15 sham images, n=14 SBR images (3 biological replicates). Day 70: n=9 sham images, n =10 SBR images (2 biological replicates). **B)** Projection of *Creb3l3* and *Klf4* transcript enrichment onto the UMAP plot shows increased expression with enterocyte maturation. Color scale bar indicates relative intensity of gene expression. **C)** Violin plots showing differential expression of *Creb3l3* and *Klf4* between sham and SBR. **D)** The length down from the villus tip was measured for appreciably higher *Creb3l3* (left, 3 sham and 3 SBR biological replicates) and Adenosine deaminase (ADA, right, 3 sham and 4 SBR biological replicates) expression, and represented as a percent of total villus length. Representative IHC staining for ADA in sham and SBR mice at post-operative day 7 is shown. Images are at 20X, scale bar = 100 μm. **E)** Relative *Creb3l3* expression in SI from day 3 post-operative sham and SBR mice (n=3 sham and n=5 SBR mice) was measured using qPCR. **F)** Interactome of genes upregulated in SBR epithelium. Genes in red are involved in RA signaling. All graphs are presented as mean ± SD. ns= not significant, *= *p*<0.05, **= *p*<0.01, ***= *p*<0.001, ****=p<0.0001.

Visualization of *Creb3l3* expression via RNA-FISH also provided valuable information on the localization of its expression within the SI epithelium. According to a recent study, 83% of enterocyte genes demonstrate spatial zonation during homeostasis, where *Creb3l3* and *Klf4* transcripts were found to localize to the upper villus^13^. This is consistent with our UMAP analysis, which showed increased *Creb3l3* and *Klf4* expression as enterocytes mature and migrate toward the villus tip (**Figures 2a, 2c, and 6b, 6c**). The earlier study also reported that upper villi are collectively enriched in transcripts associated with lipoprotein biosynthetic processes, including SBR enriched transcripts *Apoa4* and *Apoc3*^13^. Given the relative expansion of mature proximal enterocytes as a percent total epithelium in SBR, the associated enrichment of lipid processing transcripts, and the enrichment in mature proximal enterocyte transcription factor *Creb3l3* within SBR cells, we hypothesized that RNA-FISH would reveal a relative expansion of *Creb3l3* expressing cells along the villus axis after SBR. Indeed, sham mice exhibited *Creb3l3* expression along 50.7% ± 3.9% of their villi, vs 94.1 ± 1% in SBR (*p*< 0.01, **Figure 6d**), suggesting preferential expansion of enterocytes with transcriptional profiles typically found in villus tip cells, following SBR. To explore this further, we performed immunohistochemistry for an additional villus tip marker, adenosine deaminase (ADA)^13^. This confirmed increased relative concentration and distribution of ADA in the upper villi of SBR mice (22.4% ± 3.5% of upper villi with relatively intense expression in sham, vs 34.7% ± 3% in SBR, *p*< 0.05) (**Fig. 6d**). Together, these results led us to conclude that stereotypical villus axis zonation patterns are at least partially abrogated during adaptive challenge, likely underscoring the expanded mature enterocyte population in SBR.

### Interactome analysis indicates regional reprogramming is driven by Retinoic Acid (RA) signaling

Although *Creb3l3* was elevated at day 7 and sustained through day 70 after SBR—suggesting a continued role in maintaining adaptation—no significant differences in *Creb3l3* expression were observed at day 3 post-surgery (**Figure 6e**). This suggested that inductive upstream signaling at earlier stages of adaptation may be critical to driving proximalization, which is subsequently mediated and sustained by *Creb3l3*. To investigate this, we generated *in silico* interactomes from single-cell gene expression profiles of all analyzed sham and SBR cells to determine which differentially expressed genes were most strongly co-expressed, thereby inferring gene-gene relationships and pathways (**Figure 6f**). This approach identified a network of interacting genes induced by SBR, including *Sepp1*, *Apoa1*, *Fabp2*, and *Rbp2*, the upregulated expression of which we confirmed above (**Figures 4 & 5**). We used a total of 59 genes from the SBR interactome to perform gene list functional enrichment analysis (5 genes from the interactome were excluded from analysis as they were absent from the database)^31^. This analysis generated a list of pathways, including “Lipid digestion, mobilization, and transport” (*p*=2.290E-11) and “Digestion of dietary carbohydrate” (*p*=2.857E-8), in addition to “PPAR Signaling Pathway” (*p*=3.933E-7), and “Retinoid metabolism and transport” (*p*=1.935E-8).

In the context of this study, retinoid metabolism was of particular interest since it has been shown to play a key role in structural adaptation^21–23, 32, 33^. Retinoids are derived from vitamin A, which must be obtained from the diet^34^, and are mostly absorbed in proximal SI^35^. Retinoic acid (RA) is the intracellularly bioactive hydrolysis derivative of vitamin A, and it interacts with retinoid X receptor (RXR) and retinoic acid receptor (RAR) heterodimers, which bind to retinoic acid response elements (RAREs) within the nucleus, to drive effects^36^. Notably, mice deficient in dietary Vitamin A do not adapt after SBR, and it was observed that RA drives adaptation in part via regulation of enterocyte proliferation, migration, and apoptosis^32, 33^. However, relatively little detail on the molecular changes induced by RA has been revealed so far.

To investigate how RA induces the transcriptional changes accompanying SBR, we mined our dataset for genes differentially expressed between sham and SBR treatments which were either putative targets of RA signaling based on the literature (**Table 2**), or contain a predicted RARE as determined by FIMO Motif Search^37^ (**Table 3**). As a result, we found 45 genes differentially expressed between sham and SBR that likely respond to RA signaling (**Table 2**). Several of these genes including *Plb1* (phospholipase B1), *Rbp2* (retinol binding protein 2), *Apoa1* (apolipoprotein A1), *Apoa4* (apolipoprotein A4), *Apob* (apolipoprotein B), and *Apoc3* (apolipoprotein C3) (highlighted in red in **Fig 6f**), are active in retinoid metabolism and transport, four of which were are also found in our list of the top 10 most differentially expressed genes in SBR (**Table 1**). Importantly, *Rbp2* (1.1 log fold change enriched, *p*<0.001) is preferentially induced by RA in differentiated cells^38^, such as the expanding enterocyte population we observe after SBR. Furthermore, motif analysis indicated that *Creb3l3*, our main proximal transcription factor, contains a RARE and can be activated by RXRα based on ENCODE transcription factor targets (**Table 3**). *Klf4*, an additional transcription factor identified in our dataset, is also influenced by RA signaling through RARα^39^ (**Table 2**). Together, these findings confirm that RA signaling is induced in cells responding to SBR, placing these signals upstream of the key transcriptional changes we observe, supporting a crucial role for RA signaling in adaptation.

**Table 2.**
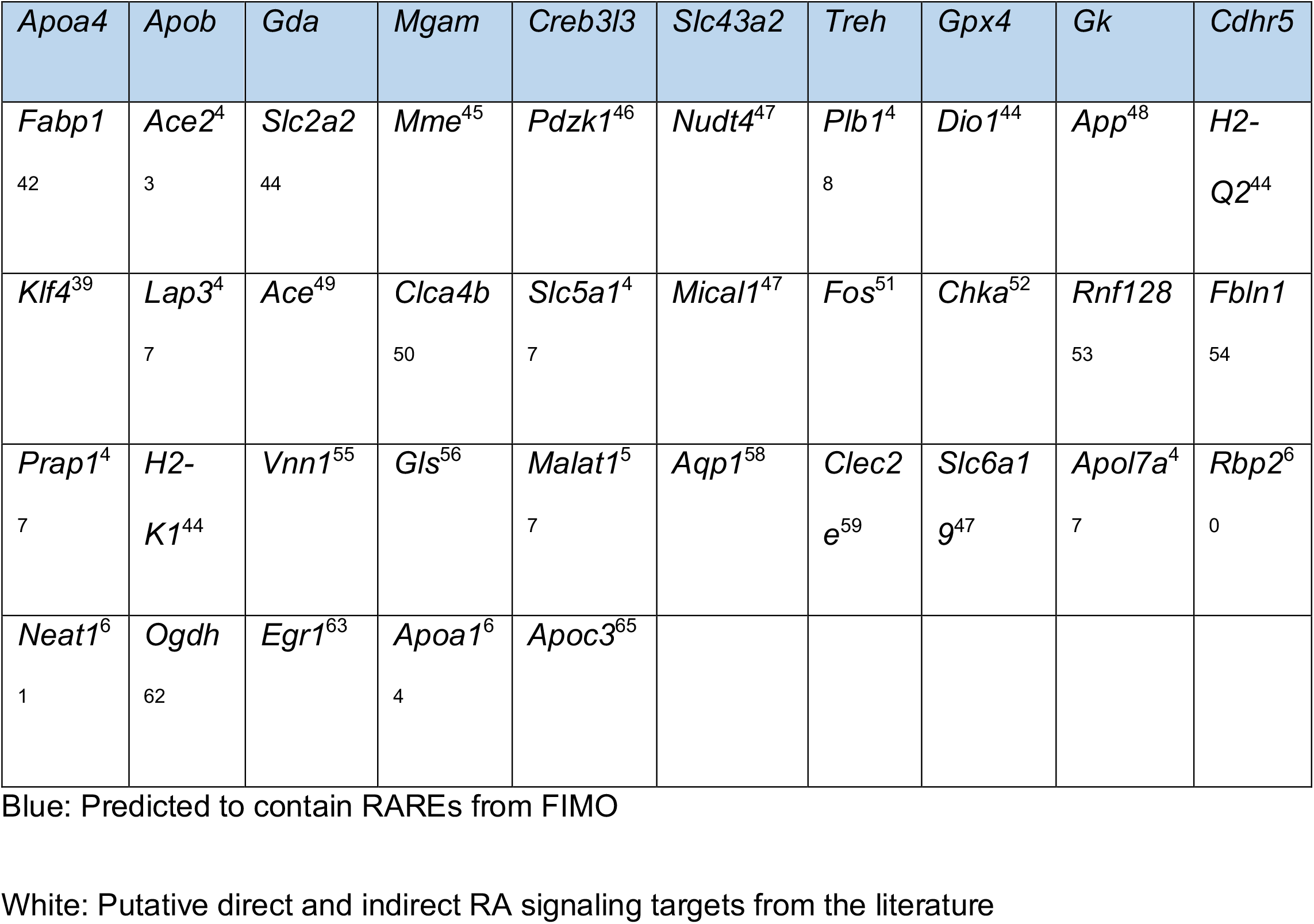
Genes increased in SBR vs sham that are Putative Responders to RA Signaling

**Table 3.** . Genes identified in SBR vs sham that are target genes of RXR*α* from ENCODE Transcription factor targets dataset (human)

|  |  |  |  |  |  |  |  |  |  |
| --- | --- | --- | --- | --- | --- | --- | --- | --- | --- |
| *Fabp1* | *Apoc3* | *Sepp1* | *Leap2* | *Ephx2* | *Apob* | *Apoa1* | *Ace2* | *Slc2a2* | *Dnase1* |
| *Pdzk1* | *Creb3l3* | *Slc43a2* | *Acsl5* | *Pls1* | *Chka* | *Ano6* | *Rnf128* | *Fbln1* | *Prap1* |
| *Rfk* | *Khk* | *Malat1* | *Pepd* | *Gsdmd* | *Neat1* | *Ogdh* | *Dhrs1* | *Gpx4* | *Egr1* |
| *Gk* | *Acox1* | *Cdhr5* | *Nudt4* |  |  |  |  |  |  |

http://amp.pharm.mssm.edu/Harmonizome/gene_set/RXRA/ENCODE+Transcription+Factor+Targets

The results presented here so far, together with previous findings, support a model in which RA signaling induces expression of proximal SI transcriptional programs, mediated by *Creb3l3* and possibly *Klf4*, to drive regional reprogramming as an adaptive response. As RA is a hydrolysis derivative of vitamin A, we next investigated potential RA signaling sources, and did not find differential expression of RA biogenesis genes between sham and SBR populations. This would suggest SBR epithelium does not upregulate RA biogenesis genes to increase RA signaling but, rather, may be responding to increased vitamin A exposure after SBR. This is consistent with a prior report showing epithelial RA production is dependent on substrate concentration, implicating an “active saturable enzymatic conversion process”^40^. It is also consistent with a previous study which demonstrated decreasing concentrations of vitamin A derivatives from proximal to distal SI tissue, suggesting that a gradient of bioavailability may impact the regional effects of these compounds within the SI^41^. Based on these observations, we propose a simple model, where after SBR, the ileum becomes exposed to dietary stimuli, including vitamin A, which previously would have been processed in the resected, more proximal, SI. We thus hypothesize that SBR disrupts the endogenous RA gradient of the small intestine, thereby promoting RARE-mediated transcription in ileal enterocytes due to novel exposure to RA.

## Discussion

Here, we report epithelial single-cell analysis of ileum following proximal SBR, showing expansion of cells identified as mature proximal enterocytes in SBR vs. sham mice. SBR enterocytes differentiated significantly on the basis of solute and nutrient transporters typically associated with proximal SI, especially with regard to lipid metabolism. Because the duodenum and jejunum absorb the majority of nutrients under normal conditions, we propose that this “regional reprogramming,” driven by transcriptional proximalization, is a critical component of the adaptation response to SBR. This is a novel principle, as the typically studied structural adaptation, while increasing absorptive surface area, does not necessarily facilitate functional adaptation^5^. Rather than simple tissue hyperplasia, as others have suggested^66^, we demonstrate that enterocyte-level alterations in transcriptional profiles occurs after SBR, in order to mimic proximal SI function. These findings highlight the significant contribution that single-cell analysis makes toward our understanding of organ pathophysiology.

While investigating causative signaling mechanisms driving these changes, we identified upregulation of two transcription factors associated with proximal SI: *Creb3l3* and *Klf4*. Furthermore, upstream analysis reiterated the importance of retinoid metabolism to the adaptation response, and the depth of our analysis allowed identification of previously undescribed transcriptional changes likely mediated by RA after SBR, including *Creb3l3* and *Klf4*. We, therefore, conclude that a critical component of the adaptation response to massive proximal SBR is transcriptomic “proximalization” of distal SI enterocytes, and that this is driven at least in part by RA signaling which is upstream of “proximalization” transcription factors and signaling cascades.

RA is derived entirely from the diet. Following proximal SBR, the ileum becomes exposed to nutrients in the luminal content which would otherwise have been largely absorbed more proximally, including Vitamin A^35^. A series of studies utilizing an ileostomy model of SGS in mice and zebrafish have shown the critical effects of mechanoluminal flow on the structural adaptation process after SBR, including loss of structural adaptation and cellular proliferation in the distal bowel when isolated from the flow of luminal contents^67, 68^. The importance of luminal nutrition or enteral feeding has also been highlighted by other groups^69, 70^. As such, exposure to increased levels of dietary Vitamin A is a likely mechanism driving *structural* adaptation, consistent with prior reports^24–26, 33, 3^. In the current study, we have identified novel regulatory networks and transcriptional changes downstream of RA signaling, which likely drive *regional reprogramming* as well.

In addition to Vitamin A, the ileum also becomes exposed to higher levels of dietary fatty acids after SBR, and this likely constitutes an additional driving force for adaptation. In line with this, our lab has previously demonstrated that a high-fat diet enhances villus growth following SBR, but not sham surgery, though enhanced structural adaptation did not correlate with enhanced functional adaptation (weight gain), despite increases in fatty acid transporters such as CD36^71^. At the same time, this study did not examine the transcripts found to be most significant in the current dataset (**Table 1**), and it may be that ligands designed to induce those specific transcripts could have a different effect. Regardless, these findings again demonstrate the potential for incongruence between structural and functional adaptation, indicating the adaptation process is inherently multifaceted, and different aspects likely rely on different stimuli. For example, two of the upregulated transcripts in SBR— *Fabp1* and *Fabp2* (**Table 1**)— are regulated independently of one another, with *Fabp1* activated by peroxisome proliferator-activated receptor alpha (or PPARα), in response to dietary fatty acids, and *Fabp2* hormonally by PYY^72^.

The PPAR signaling pathway was implicated by our interactome analysis. We investigated this further since PPARs form heterodimers with RXRs to activate PPAR response elements (PPREs) in the induction regions of many genes involved in lipid metabolism^73, 74^, including *Creb3l3*^75^. Interestingly, it is thought the ratio of RA binding protein to fatty acid gene expression determines the functional outcome of RA signaling^76^ and, further, treatment of mice with a PPARα agonist-induced villus growth by facilitating cell differentiation^77^, similar to the adaptation phenotype observed after SBR in which there are elongated villi with a preponderance of mature enterocytes. Ultimately, we did not observe changes in *Pparα* expression (1.5% decrease 3 days after SBR via qPCR, *p*=0.96, not shown), nor did we observe significant changes in *Pparδ* expression (55.2% decrease 3 days after SBR, *p*=0.16, not shown). Less is known about PPARδ, though it is thought to interact with corepressors and function as an inhibitor of PPARα^78^. Given these findings, we suspect that either 1) PPAR signaling is important to SBR adaptation, but was not captured in our analysis temporally, or 2) minimal to no transcriptional change in these specific genes is needed to drive a significant biological effect.

Finally, another key finding from this study was the abrogation of villus zonation patterns during adaptive challenge, with the expansion of villus tip transcript *Creb3l3* and ADA along the lengths of villi. The upper villus is typically responsible for fatty acid absorption, which is less metabolically demanding, and so preferential fat absorption/chylomicron secretion at the upper villus parallels the decreased bioavailability of oxygen in this area^13^. Indeed, we have previously demonstrated that adaptation to SBR is associated with relative hypoxia^79^, and so the metabolic incentive to prioritize fatty acid absorption is at least two-fold after SBR: starvation and relative hypoxia. Mechanisms underlying redistribution of villus zonation during physiologic challenge warrant further investigation.

In summary, our analysis has revealed a significant shift in metabolic machinery and regional identity at the enterocyte level following SBR. Moving forward, additional studies are warranted to better delineate causal factors driving changes between sham and SBR enterocytes, in conjunction with or independent of RA signaling. This is especially true considering prior studies that demonstrated proliferative and morphometric effects of circulating factors on structural “jejunalization” of ileum following SBR^80, 81^, which implicates non-luminal stimuli in driving structural adaptation. Similar studies exploring molecular changes in response to circulating factors would provide further insight. Discerning the stimuli for functional proximalization of ileum following SBR will prove critical, as it will provide insight toward targeted therapeutic approaches, via the enteral or parenteral route, for patients suffering from SGS. Targeted therapeutic approaches combining aspects of adaptation from a structural and molecular avenue could induce heightened adaptive responses, yielding better patient outcomes.

## Conclusions

Here, we have characterized the transcriptome of adapted intestinal epithelium at the single-cell level following massive SBR, a laboratory model for SGS, using scRNA-seq. Our analysis revealed the emergence of unique enterocyte gene expression patterns between sham and SBR mice, which distinguished themselves on the basis of proximal vs distal SI patterning and cell identity, including critical absorptive and anti-inflammatory/immunological features. Pathways driving these changes, such as RA signaling, deserve further investigation, as they underlie the functional aspects of adaptation to SGS, which allow progressive tolerance of enteral feeding and weaning from parenteral nutrition.

## Methods

### Mice

50% proximal SBR was performed on male C57/B6 mice at 8-12 weeks of age, according to standard protocol^3^. Briefly, the SI was extruded via a midline laparotomy and the ileocecal valve (ICV) identified. The SI was transected 12cm proximal to the ICV, and ∼ 2cm distal to the ligament of Treitz. The intervening SI was removed, the mesentery ligated with 3-0 silk suture, and the proximal and distal ends approximated with interrupted 9-0 nylon stitches. Sham surgery, consisting of distal transection and anastomosis only, was performed as control. Peritoneum and skin were approximated in separate layers, animals were resuscitated with a subcutaneous bolus of normal saline (repeated on post-operative day 1) and co-housed in a 33°C incubator until the end of the 7 day experiment. For longer studies (70 days), mice were moved to room temperature at day 7. Liquid diet (LD; PMI Micro-Stabilized Rodent Liquid Diet LD 101; TestDiet, St. Louis MO) was initiated 24h prior to surgery, withheld the morning of surgery, and subsequently provided on post-operative day 1 until the end of the experiment. Food and water were available *ad libitum*, and animals were housed under 12h light darkness cycles with corn cob bedding and nestlet enrichment. All surgical and animal care procedures were approved by the Washington University Institutional Animal Care and Use Committee, and meet Animal Research: Reporting of In Vivo Experiments standards.

### Tissue isolation and processing

At day 7 after surgery, epithelium was isolated from a 1cm segment of SI 3cm distal to the anastomosis, similar to previously published protocols^82, 83^. Briefly, the SI was flushed with ice cold sterile saline, filleted lengthwise, and placed in a conical tube containing ice cold 30 mM EDTA in phosphate buffered saline (PBS). After 15 minutes on ice without agitation, the SI segment was transferred to a fresh conical tube containing 30 mM EDTA in PBS, briefly shaken, and placed in a 37°C water bath for 15 min. Subsequently, the tube was shaken aggressively by hand for 2 minutes. Sub-epithelial tissue floated to the top and was removed, and epithelium was pelleted by centrifugation. Epithelium was then re-suspended in a 0.3U/ml dispase solution (07923, Stem Cell Technologies; Cambridge, MA) and incubated at 37°C for 15 min, shaking every 2 minutes. The solution was then quenched with media containing fetal bovine serum to a final concentration of 5%, pipetted several times, and sequentially passed through 100, 70, and 40μm filters. Single-cell suspensions were confirmed by microscopy, pelleted by centrifugation, and resuspended in 200 μl ice cold PBS. 800 μl ice cold 100% methanol was then added, dropwise, with gentle mixing between drops. Samples were immediately stored in 80% methanol in PBS at −80°C, according to Alles *et al*. 2017^84^, for later processing. For Western Blotting analysis, whole epithelium was isolated using 30 mM EDTA, pelleted by centrifugation, and lysed in sodium dodecyl sulfate sample buffer (50 mmol/L Tris-HCL, pH 6.8, 2% sodium dodecyl sulfate, 10% glycerol, and 5% 2-mercaptoethanol). Lysate was heated to 100°C and stored at −20°C prior to processing. Protein concentration was measured using the RC DC (reducing agent and detergent compatible) Protein Assay Kit II (5000122, Bio-Rad; Hercules, CA). For qPCR experiments, RNA was isolated from homogenized whole SI using the standard Trizol method. RNA concentration was measured using a NanoDrop Spectrophotometer (ND- 1000, NanoDrop Technologies; Wilmington, DE). 1μg RNA was converted to cDNA using qScript cDNA Synthesis Kit (95047, Quanta Bio, Beverly, MA), according to the manufacturer’s instructions, and stored at −20°C until use.

### scRNA-seq library preparation

For single-cell library preparation on the 10x Genomics Chromium platform, we used: the Chromium Single 3′ Library & Gel Bead Kit v2 (PN-120237), Chromium Single Cell 3′ Chip kit v2 (PN-120236) and Chromium i7 Multiplex Kit (PN-120262), according to the manufacturer’s instructions in the Chromium Single Cell 3′ Reagents Kits V2 User Guide. Methanol-fixed cells from sham (n=3) and SBR (n=3) animals were pooled for the first batch. Methanol-fixed cells from sham (n=2) vs SBR (n=1) were processed individually in a separate experiment for a final sample size of sham (n=5) and SBR (n=4). Just prior to cell capture, methanol-fixed cells were placed on ice, then spun at 3000rpm for 5 minutes at 4°C, followed by resuspension and rehydration in PBS, as previously described^84^. Resulting cDNA libraries were quantified on an Agilent Tapestation and sequenced on an Illumina HiSeq 2500.

### scRNA-seq analysis

The Cell Ranger v2.1.0 pipeline was used to align reads to the mm10 genome build, and generate a digital gene expression (DGE) matrix: (https://support.10xgenomics.com/single-cell-gene-expression/software/downloads/latest). For initial filtering of these DGE matrices, we first excluded cells with a low number (<200) of unique detected genes. We then excluded cells for which the total number of unique molecules (UMIs, after log10 transformation) was not within three standard deviations of the mean. This was followed by the exclusion of outlying cells with an unusually high or low number of UMIs/genes given their number of reads by fitting a loess curve (span = 0.5, degree = 2) to the number of UMIs/genes with number of reads as predictor (after log10 transformation), removing cells with a residual more than three standard deviations the mean. Finally, we excluded cells in which the proportion of the UMI count attributable to mitochondrial genes was greater than 25%. Raw and processed data files are available via GEO: accession number GSE130113. After filtering and normalization of the DGE, the R package Seurat^7^ (Version 3) was used to cluster and analyze the single-cell transcriptomes. Independent biological replicates from the sham and SBR surgeries were integrated by Canonical Correlation Analysis (CCA), identifying common sources of variation to align the datasets, reducing batch effects^8^. Highly variable genes were identified and used as input for dimensionality reduction via CCA. The resulting CCs and the correlated genes were examined to determine the number of components to include in downstream analysis, followed by clustering and visualization via (Uniform Manifold Approximation and Projection) UMAP^9^.

### Quadratic Programming analysis to assess cell identity and state

Quadratic programming (QP), previously described in Treutlein *et al*. 2016^10^ and successfully modified and used by our group in Biddy *et al.* 2018^12^, was used to score cell identity. Here, we created a reference of SI epithelial cell types, collected previously (from Haber *et al*. 2017^11^). The R Package, QuadProg, was used for QP to generate cell identity scores, modifying our earlier approach by modeling cell type classification as a multivariate linear regression, solved for fractional cell types or identities. This approach enables cell identities to be categorized. In addition, it enables more subtle changes in cell identity to be quantified.

### Immunohistochemistry

Ileal tissue adjacent to the region collected for single-cell preparation was fixed in 10% neutral buffered formalin, paraffin embedded, and sectioned at a thickness of 5 μm. Deparaffinization and immunolocalization were performed as previously described^85^. Briefly, slides were deparaffinized in xylene, rehydrated in sequential ethanol baths, and prepared in 3% hydrogen peroxide in methanol. Antigen retrieval was performed using 1x Diva Decloaking Solution (DV2004, Biocare Medical, Pacheco, CA), and blocking was performed using the Avidin-Biotin kit (AB972L, Biocare Medical). Primary antibodies were diluted in Da Vinci green (PD900L, Biocare Medical) and incubated overnight at 4°C. Slides were rinsed in PBST, incubated in biotin-labeled secondary IgG diluted in PBST, rinsed in PBST, incubated in streptavidin-HRP diluted in PBST, developed in DAB (D9015, Sigma, St. Louis, MO), counter stained with Hematoxylin and bluing agent, run in successive dilutions of ethanol, and xylene, and cover-slipped using MM 24^TM^ mounting medium (100109, Surgipath, Richmond, IL). Of note, samples used for confirmatory staining were from a different litter of mice than those used for scRNA-seq analysis. This was done to validate consistency of results across cage and littermates, which has been previously reported as a confounding variable in murine GI research^86^. At least 3 sham and 3 SBR samples were analyzed after surgery. 20x images representative of the sample were obtained by a blinded investigator using a Nikon Eclipse 80i with Ds-Ri2 camera and NIS Elements V4.3 software (Nikon Instruments, Inc., Melville NY).

### Western Blotting

20 μg of each sample and 15uL Novex® Sharp Pre-stained Protein Standard (57318, Invitrogen, Carlsbad, CA) was loaded onto a 18% polyacrylamide gel. Western blotting was performed on a nitrocellulose membrane (IB301001, Invitrogen) after a dry transfer using the iBLOT® Gel Transfer Device (IB1001, Invitrogen). The membrane was blocked with 5% BSA in PBST for 1 hour at room temperature, followed by overnight incubation in primary antibody at 4°C. The membrane was washed in PBST 3 times in 10-minute increments, incubated for 1 hour in secondary antibody at room temperature, washed in PBST, and developed using GE Healthcare Amersham^TM^ ECL^TM^ Western Blotting Detection Reagents (6883S, GE Healthcare, Chicago, IL). Image Lab Software (Bio-Rad) was used to detect and quantify proteins.

### Quantitative polymerase chain reaction (qPCR)

cDNA was amplified using TaqMan Gene Expression Master Mix (4369016, Applied Biosystems, Foster City, CA) and the specified primer probe on the Applied Biosystems 7500 Fast Real-Time PCR system. Primer probes were *Actb* (Mm02619580_g1, endogenous control), *Creb3l3* (Mm00520279_m1), *Ephx2* (Mm01313813_m1), and *Rbp2* (Mm00436300_m1), all from Applied Biosystems.

### Immunofluorescence

Ileal tissue adjacent to the region collected for single cell-preparation was fixed overnight in 4% paraformaldehyde and then overnight in 30% sucrose before being embedded in O.C.T. Compound (23-730-571, Fischer Healthcare, Houston, TX), sectioned at 5 μm, and stored at −80°C until use. Slides were washed in PBS and blocked in 5% goat serum with 0.3% Triton X-100 (T8787, Sigma) prior to incubation with primary antibody in antibody staining solution (1% goat serum and 0.3% Triton X-100) at 4°C overnight. Slides were rinsed in PBS for 5 minutes three times, and secondary antibody was applied in antibody staining solution for 1 hour at room temperature. 300nM DAPI (D1306, Invitrogen, Lot 1802085) was applied for 1 minute, and slides were rinsed in PBS for 5 minutes three times before a cover slip was applied using ProLong Gold Antifade Mountant (P10144, Invitrogen). 40x images representative of the sample were obtained by a blinded investigator using a Nikon Eclipse 80i with Ds-Ri2 camera and NIS Elements V4.3 software (Nikon Instruments, Inc., Melville NY). Images were subsequently analyzed using ImageJ (National Institutes of Health, Bethesda, MD) for FABP6 and SEPP1, or unbiased computational quantification (FABP1, see “Quantitative analysis of RNA-FISH images”). Representative images were chosen.

### ImageJ Quantitative Analysis

Briefly, for SEPP1, regions of interest were drawn tracing the edges of all villi in the image. Each image was thresholded to the same intensity and the percent area within regions of interest expressing fluorescence was collected for each technical replicate. For FABP6, the full area of each villus was chosen as a region of interest. The mean intensity of that region was calculated to represent the fluorescence of the diffuse FABP6 staining. For percent villus occupancy, ImageJ was used to measure the areas of *Creb3l3* expression or ADA expression, and this was divided by the total villus length. For *Creb3l3*, there were n=12 sham villi (3 biological replicates) and n=11 SBR villi (3 biological replicates). For ADA, there were n=12 sham villi (3 biological replicates) and n= 16 SBR villi (4 biological replicates).

### RNA fluorescent *in situ* hybridization (RNA-FISH)

Tissues were fixed and sectioned as previously described in methods for immunohistochemistry and immunofluorescence, above. RNA-FISH was performed using the RNAscopeⓇ Multiplex Fluorescent v2 kit (323100, Advanced Cell Diagnostics, Newark, CA), following the protocol for Fixed Frozen Tissue. Briefly, tissue was pre-treated with RNAscopeⓇ Hydrogen Peroxide (322335, Advanced Cell Diagnostics), target retrieval was performed for 5 minutes, and tissue was treated with RNAscopeⓇ Protease III (322337, Advanced Cell Diagnostics). Then, specified probes were hybridized using the RNAscopeⓇ HybEZ II Oven (321710/321720, Advanced Cell Diagnostics). Probes were then amplified and the HRP signal was developed using TSA Fluorescein Plus Evaluation Kit (NEL741E001KT, Perkin Elmer, Waltham, MA). Finally, slides were counterstained with DAPI (323108, Advanced Cell Diagnostics) and mounted with ProLong Gold antifade reagent (P10144, Life Technologies Corporation, Eugene, OR). Imaging was performed on an Olympus FV1200 Confocal Microscope, where multiple 60x images were taken per sample of which 2-3 samples were used for each timepoint (day 7 vs. day 70) in each treatment (sham vs. SBR). Images were subsequently analyzed using unbiased computational quantification and representative images were chosen.

### Quantitative analysis of RNA-FISH images

RNA-FISH images were processed with a custom python script to quantify gene expression level at single-cell resolution. Individual cell segmentation was achieved based on the nuclear signal. First, DAPI images were transformed into binary images by thresholding the fluorescent signal. The threshold values were determined by the Otsu method^87^. Binarized nuclei images were processed by the watershed segmentation method to completely separate individual objects. The images were subjected to 2-step quality check: filtering of objects and filtering of images. First, inappropriately-sized objects were filtered to remove noise and cell multiplets. Then, images with a large number of inappropriate objects were removed. The intensity of the fluorescent signal per individual cell area was then quantified. Fluorescent signals per image were averaged to obtain mean signals per sample and treatment.

### Gene Co-Expression and Interactome Analysis

To further reveal relationships among differentially expressed genes following SBR, we constructed gene co-expression networks using weighted gene correlation network analysis (WGCNA), adapted for single-cell analysis as in https://hms-dbmi.github.io/scw/WGCNA.html. The analysis was performed using the R package, WGCNA. In brief, differentially expressed genes between SBR and sham samples were identified via Seurat analysis, as above (Using the function, *FindMarkers*). Genes that were upregulated in SBR treatment were used for network construction. Correlations of each gene pair among all significantly differentially expressed genes were then calculated. A gene-gene correlation matrix was used to construct an adjacency matrix by raising the correlations to a soft-threshold power, from which Topological Overlap Matrix (TOM) was further computed to remove spurious correlations. With the TOM matrix, the algorithm identifies modules/clusters of genes via clustering using Ward’s method and Dynamic Branch Cut methods. Here, to identify the most significant connections, we select the top 5% of the most differentially expressed genes for visualization. The network was visualized using CytoScape based on the TOM matrix.

### Statistical analysis

Statistical comparison of the QP-generated identity scores between the two groups, sham and SBR, was performed using an unpaired Student’s *t*-test. For Western Blot, qPCR, and percent villus occupancy studies, statistical analyses were performed using Prism 7.0 (GraphPad Software; La Jolla, CA). Differences in protein expression and mRNA expression between sham and SBR were assessed using unpaired Student’s *t*-tests. Graphs with error bars represent mean + /- SD. *P*<0.05 was considered significant. For immunofluorescence quantitative analysis using ImageJ, SEPP1 and FABP6 replicate values were averaged between sham and SBR treatment identities. A Wilcoxon rank sum test was performed to determine statistical significance. For quantitative analysis of RNA-FISH images, statistics were calculated using the Wilcoxon rank sum test. These processes were run with Python 3.6.7 and its libraries: scikit-image 0.13.1, numpy 1.14.3, pandas 0.24.1, oiffile 2019.1.1, matplotlib 3.0.3, seaborn 0.8.1, jupyter 1.0.0.

**Table 1.**
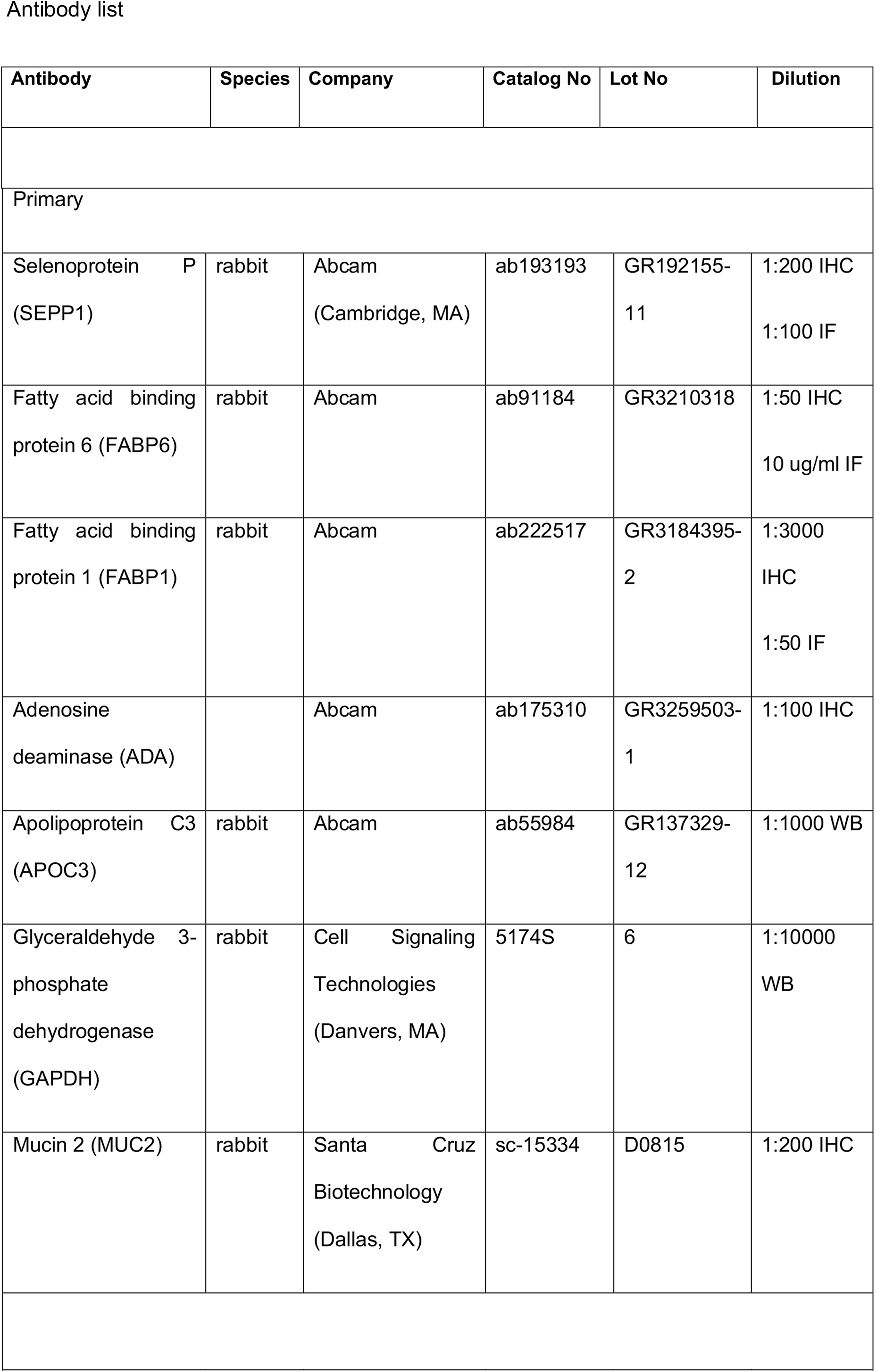
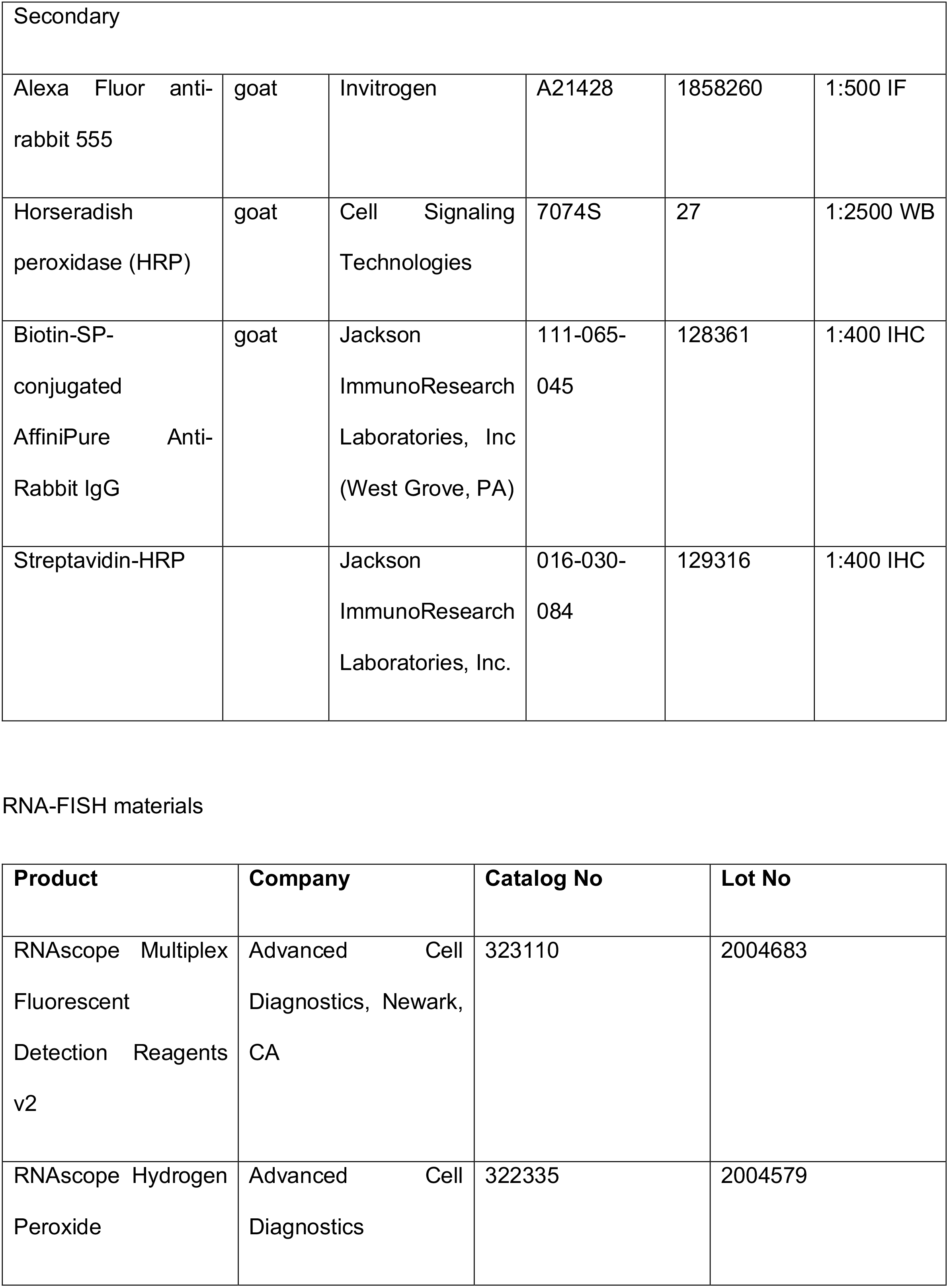
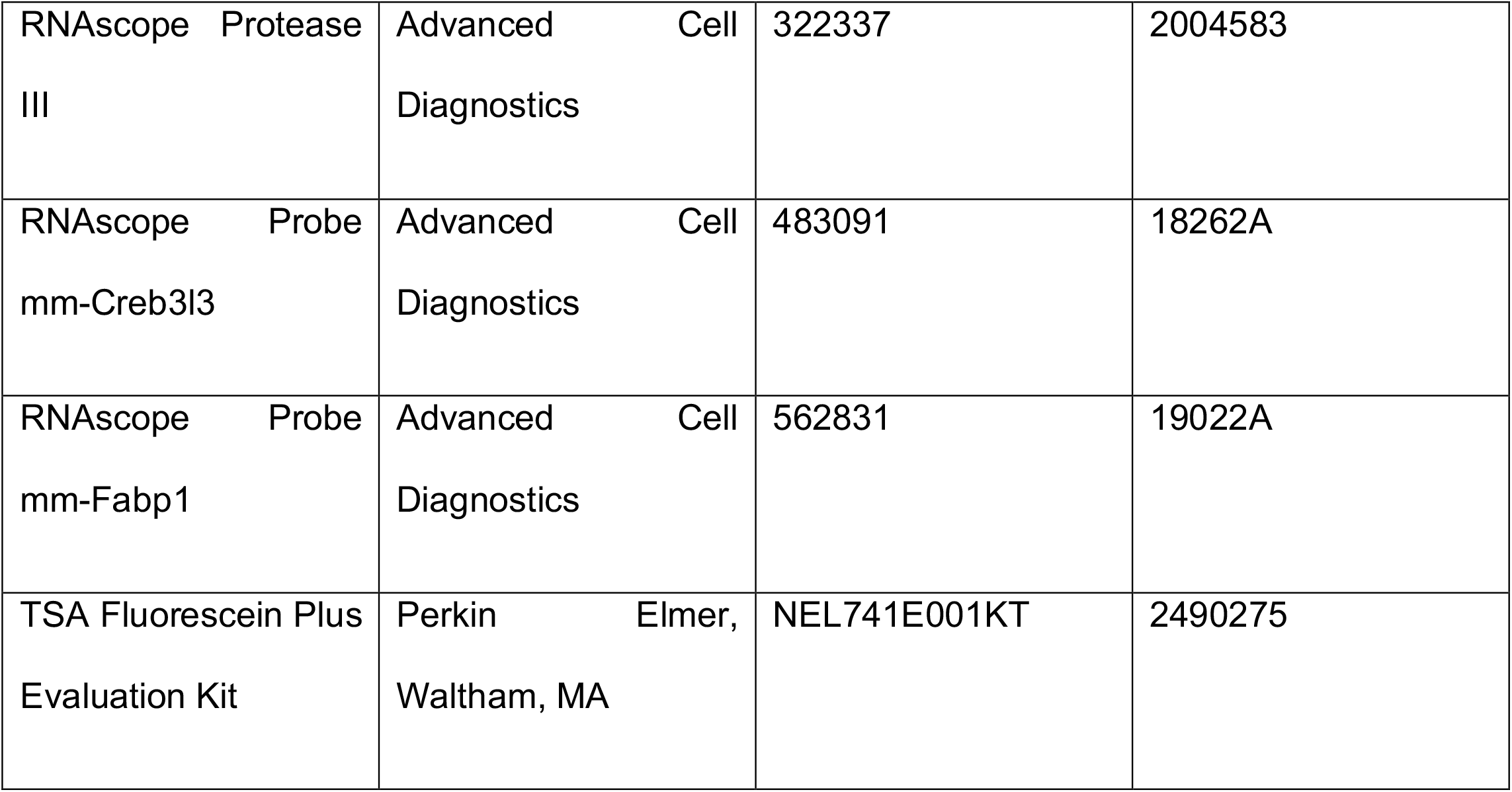
Materials.

| Antibody | Species | Company | Catalog No | Lot No | Dilution |
| --- | --- | --- | --- | --- | --- |
| Primary |  |  |  |  |  |
| Selenoprotein P (SEPP1) | rabbit | Abcam (Cambridge, MA) | ab193193 | GR192155-11 | 1:200 IHC<br>1:100 IF |
| Fatty acid binding protein 6 (FABP6) | rabbit | Abcam | ab91184 | GR3210318 | 1:50 IHC<br>10 ug/ml IF |
| Fatty acid binding protein 1 (FABP1) | rabbit | Abcam | ab222517 | GR3184395-2 | 1:3000 IHC<br>1:50 IF |
| Adenosine deaminase (ADA) |  | Abcam | ab175310 | GR3259503-1 | 1:100 IHC |
| Apolipoprotein C3 (APOC3) | rabbit | Abcam | ab55984 | GR137329-12 | 1:1000 WB |
| Glyceraldehyde 3-phosphate dehydrogenase (GAPDH) | rabbit | Cell Signaling Technologies (Danvers, MA) | 5174S | 6 | 1:10000 WB |
| Mucin 2 (MUC2) | rabbit | Santa Cruz Biotechnology (Dallas, TX) | sc-15334 | D0815 | 1:200 IHC |
| Secondary |  |  |  |  |  |
| Alexa Fluor anti-rabbit 555 | goat | Invitrogen | A21428 | 1858260 | 1:500 IF |
| Horseradish peroxidase (HRP) | goat | Cell Signaling Technologies | 7074S | 27 | 1:2500 WB |
| Biotin-SP-conjugated AffiniPure Anti-Rabbit IgG | goat | Jackson ImmunoResearch Laboratories, Inc (West Grove, PA) | 111-065-045 | 128361 | 1:400 IHC |
| Streptavidin-HRP |  | Jackson ImmunoResearch Laboratories, Inc. | 016-030-084 | 129316 | 1:400 IHC |

RNA-FISH materials
| Product | Company | Catalog No | Lot No |
| --- | --- | --- | --- |
| RNAscope Multiplex Fluorescent Detection Reagents v2 | Advanced Cell Diagnostics, Newark, CA | 323110 | 2004683 |
| RNAscope Hydrogen Peroxide | Advanced Cell Diagnostics | 322335 | 2004579 |
| RNAscope Protease<br>III | Advanced Cell<br>Diagnostics | 322337 | 2004583 |
| RNAscope Probe<br>mm-Creb3l3 | Advanced Cell<br>Diagnostics | 483091 | 18262A |
| RNAscope Probe<br>mm-Fabp1 | Advanced Cell<br>Diagnostics | 562831 | 19022A |
| TSA Fluorescein Plus<br>Evaluation Kit | Perkin Elmer,<br>Waltham, MA | NEL741E001KT | 2490275 |

## Acknowledgements

The authors would like to thank members of the Morris and Warner labs for helpful discussion. This work was supported by the Children’s Discovery Institute of Washington University in St. Louis and St. Louis Children’s Hospital MI-II-2016-544 (S.A.M.) and MI-F-2017-629 (K.M.S.); National Institutes of Health NIH grant R01-GM126112, Silicon Valley Community Foundation, Chan Zuckerberg Initiative Grant HCA2-A-1708-02799, and Washington University Digestive Diseases Research Core Center, National Institute of Diabetes and Digestive and Kidney Diseases P30DK052574 (S.A.M.); the Children’s Surgical Sciences Research Institute of the St. Louis Children’s Hospital (B.W.W.); the Association for Academic Surgery Foundation (K.M.S.). S.A.M. is supported by a Vallee Scholar Award, S.E.W is supported by National Institutes of Health 5T32GM007067-44, and K.M.S. by 4T32HD043010-14. Imaging using the Olympus FV1200 was performed using Washington University Center for Cellular Imaging (WUCCI), supported by Washington University School of Medicine, The Children’s Discovery Institute of Washington University and St. Louis Children’s Hospital (CDI-CORE-2015-505), and the Foundation for Barnes-Jewish Hospital (3770). Select histological analyses were performed with support from the Washington University Digestive Diseases Research Core Center.

## Author contributions

K.M.S.: wrote manuscript, study concept and design, obtained funding, data acquisition, analysis, interpretation, statistical analysis, figure preparation, critical review of manuscript. S.E.W.: wrote manuscript, data acquisition, analysis, interpretation, statistical analysis, figure preparation, critical review of manuscript. W.K.: analysis, statistical analysis, figure preparation. K.K.: analysis, statistical analysis. A.B., W.H.G, E.J.O., C.C., J.G.: data acquisition. B.W.W., S.A.M.: study concept and design, analysis, obtained funding, critical review of manuscript, intellectual content. S.A.M. provided overall study supervision.

